# Metagenomic insights into an enigmatic gammaproteobacterium that is important for carbon cycling in cave ecosystems worldwide

**DOI:** 10.1101/2024.08.23.608578

**Authors:** Daniel S. Jones, Katelyn M. Green, Abigail Brown, Zoë E. Havlena, Mackenzie B. Best, Diana E. Northup, Maurizio Mainiero, Benjamin Auch

**Affiliations:** New Mexico Institute of Mining and Technology, Socorro, NM, USA; National Cave and Karst Research Institute, Carlsbad, NM, USA; Biology Department, University of New Mexico Albuquerque, New Mexico, USA; Gruppo Speleologico Marchigiano, Ancona, Italy; Federazione Speleologica Marchigiana, Ancona, Italy; Phase Genomics, Seattle, WA, USA

## Abstract

Caves are windows into the subsurface through which we can directly evaluate the microbiological processes responsible for rock weathering and biogeochemical cycling in more expansive areas of Earth’s subsurface. However, many cave microbial communities are dominated by microorganisms from uncultivated groups with unknown genomic capabilities that cannot be resolved from marker gene surveys alone. An example of this is a genus of *Gammaproteobacteria* known as “wb1-P19”, which are ubiquitous and abundant in rRNA gene surveys from limestone and volcanic caves around the world. We recovered a nearly complete metagenome-assembled genome (MAG) representing a population of wb1-P19 from Lehman Caves in Great Basin National Park, Nevada, USA, and used it to identify additional MAGs representing this group from the Frasassi Caves in Italy and in publicly available databases. Although members of the wb1-P19 group have often been thought to be autotrophs that oxidize inorganic nitrogen compounds, we show that wb1-P19 are actually obligate or facultative methanotrophs capable of aerobic and anaerobic growth. Based on genomic classification, wb1-P19 are members of upland soil cluster γ (USCγ; part of the newly proposed order “*Candidatus* Methyloligotrophales”), and are likely important for methane consumption and carbon cycling in caves and other subterranean ecosystems globally.

## Main text

Caves contain robust methanotrophic microbial communities that make them globally important sinks for atmospheric methane [1-4]. Limestone caves and karst terranes also consume CO_2_, both because of carbonic acid-driven mineral dissolution, which represents a short-term CO_2_ sink [5], and sometimes because of CO_2_ uptake by autotrophic microbial communities [6]. However, many of the microorganisms in these subterranean communities are uncultivated organisms that are only known from rRNA gene surveys, and it is therefore challenging to determine their potential roles in carbon cycling and methane consumption.

One of the most ubiquitous but enigmatic members of cave microbial communities is a group of *Gammaproteobacteria* known as “wb1-P19”. This taxon was first identified in Weebubbie Cave in Australia by Holmes et al. [7], who reported a 16S rRNA gene clone that was 90.6% similar to *Thioalkalivibrio denitrificans*. Because clades of uncultivated microorganisms are often named after representative clones, “wb1-P19” was incorporated into the SILVA taxonomy [8] and now represents a genus-level group that has been found in numerous environmental samples from around the world. This taxon is frequently among the most abundant taxa in culture-independent surveys of limestone and volcanic caves [6, 9-13], sometimes more than 30% of the microbial community [14, 15], and was identified as one of three “keystone members” in a co-occurrence analysis of 128 microbial communities from karst caves in China [13]. Because they are classified as family *Nitrosococcaceae*, wb1-P19 are thought to be responsible for CO_2_ uptake and inorganic nitrogen compound oxidation (e.g., [9, 12, 16]), and have been associated with high CO_2_ uptake in carbonate mineral deposits [6]. However, inferences of its metabolism are based on rRNA gene classification alone, and it has not yet been isolated in culture or recognized in metagenomic datasets.

In a recent study on microbial communities from Lehman Caves, Nevada, USA, Havlena et al. [15] found that wb1-P19 was abundant in rRNA gene libraries in mineral deposits from an oligotrophic passage known as the Gypsum Annex. We therefore targeted samples with abundant wb1-P19 for metagenomic analysis. However, these samples had extremely low biomass, so recovering enough DNA for metagenomic library creation proved challenging. In our initial attempts, we were only able to produce one low coverage metagenomic library using Nextera library preparation (sample LMC19-11, 39 Mb). From this dataset, we recovered a 16S rRNA gene that was classified as wb1-P19, but the dataset did not have sufficient depth of coverage for binning, and we did not have enough material from this sample or library for deeper sequencing. We subsequently attempted metagenome sequencing from a different sample from the cave, which resulted in a larger library (sample LGA18-4, 2.3 Gb). From this dataset, we recovered a high coverage bin that contained a full-length 16S rRNA gene located in the middle of a 34.5 kb contig. The high coverage of this rRNA gene and other contigs in the corresponding bin indicate that the rRNA gene was correctly assigned to the MAG (208× average coverage for contigs in the MAG and 180× average coverage for the contig with the 16S rRNA gene, while other bins had 8-36× average coverage and other contigs with 16S rRNA genes were <37× average coverage). We improved the bin by assembling subsampled datasets, and after careful manual inspection and curation in anvi’o [17] (Figure S1), produced a MAG that was estimated to be 95.8% complete with 0% redundancy. Based on marker genes in the Lehman MAG, we then identified a second very similar MAG in a metagenome dataset from the Frasassi Caves that was generated using ProxiMeta Hi-C methods (Phase Genomics), as well as two closely-related MAGs in the genome taxonomy database (GTDB) [18], all of which were classified as genus “USCg-Taylor” (order and family “JACCXJ01” in the *Gammaproteobacteria*). Detailed methods are provided in the Supplementary Materials. The Lehman and Frasassi MAGs have been submitted to NCBI under Biosamples SAMN43226168 and SAMN43226270, respectively, and are also available at https://doi.org/10.6084/m9.figshare.26809915.v1.

Phylogenetic analysis of the 16S rRNA gene from the Lehman Caves MAG, and the beta subunit of the DNA-directed RNA polymerase gene (*rpoB*) from both the Lehman and Frasassi MAGs, show that the organisms represented by these MAGs do not cluster with named members of family *Nitrosococcaceae* (Figure 1). Instead, these MAGs cluster in a separate clade in the *Gammaproteobacteria*. In the *rpoB* phylogeny, the wb1-P19 cluster occurs as a sister clade to the *Nitrosococcaceae* that includes the two USCg-Taylor MAGs and some more distantly related MAGs (Figure 1a). In the 16S rRNA gene phylogeny, wb1-P19 occurs in a group with many cave sequences that are more closely related to *Thioalkalivibrio* (family *Ectothiorhodospiraceae*) than to *Nitrosococcaceae* (Figure 1b). wb1-P19 is therefore probably not a true member of either the *Nitrosococcaceae* or the *Ectothiorhodospiraceae*, but instead an unnamed group within the *Gammaproteobacteria*, consistent with the classification of USCg-Taylor as order “JACCXJ01” in the GTDB.

**Figure 1.**
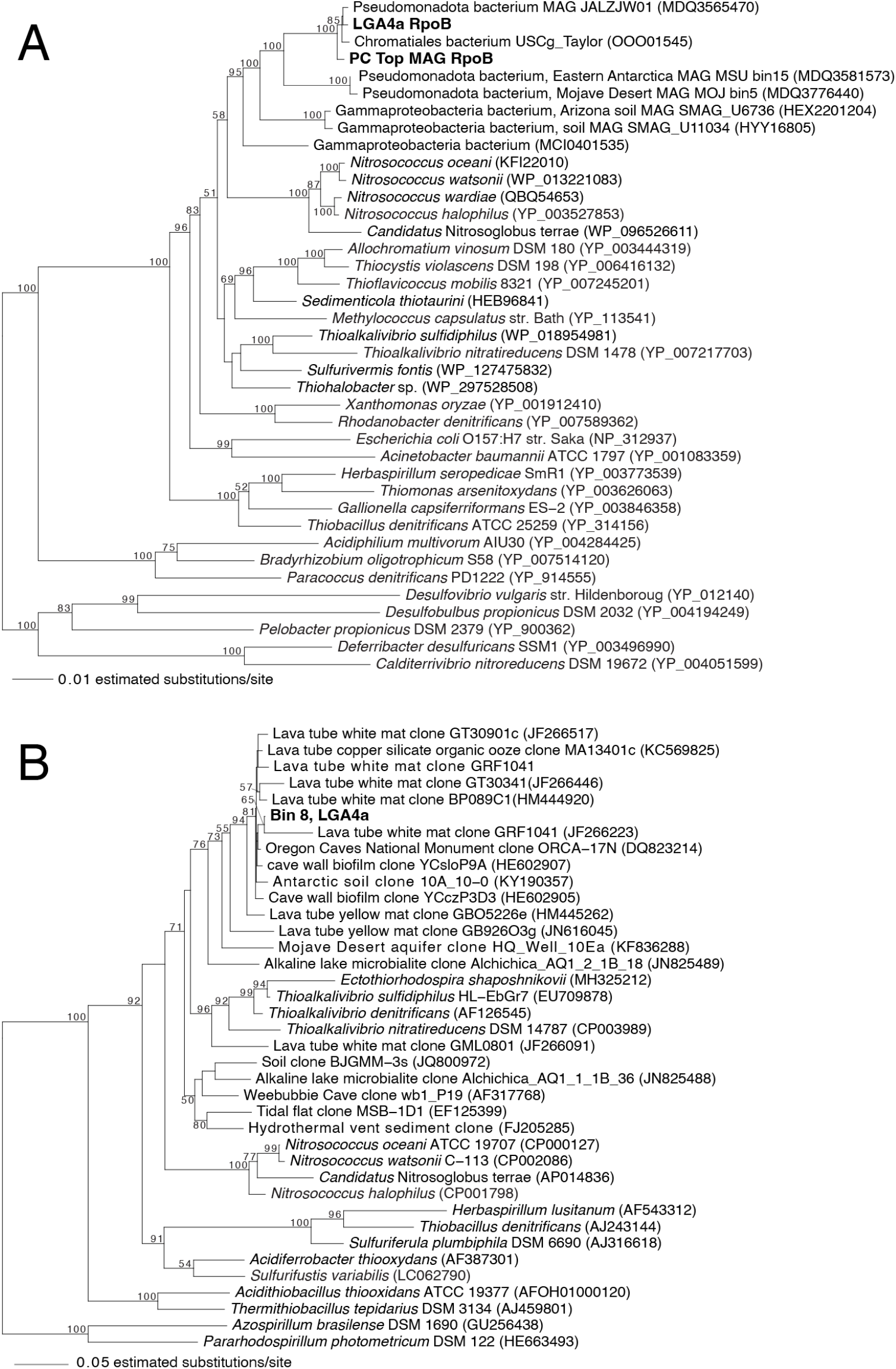
Maximum likelihood analysis of (A) *rpoB* and (B) 16S rRNA genes from the wb1-p19 MAGs from Lehman Caves and Frasassi. Numbers at nodes indicate bootstrap support ≥50.

**Figure 2.**
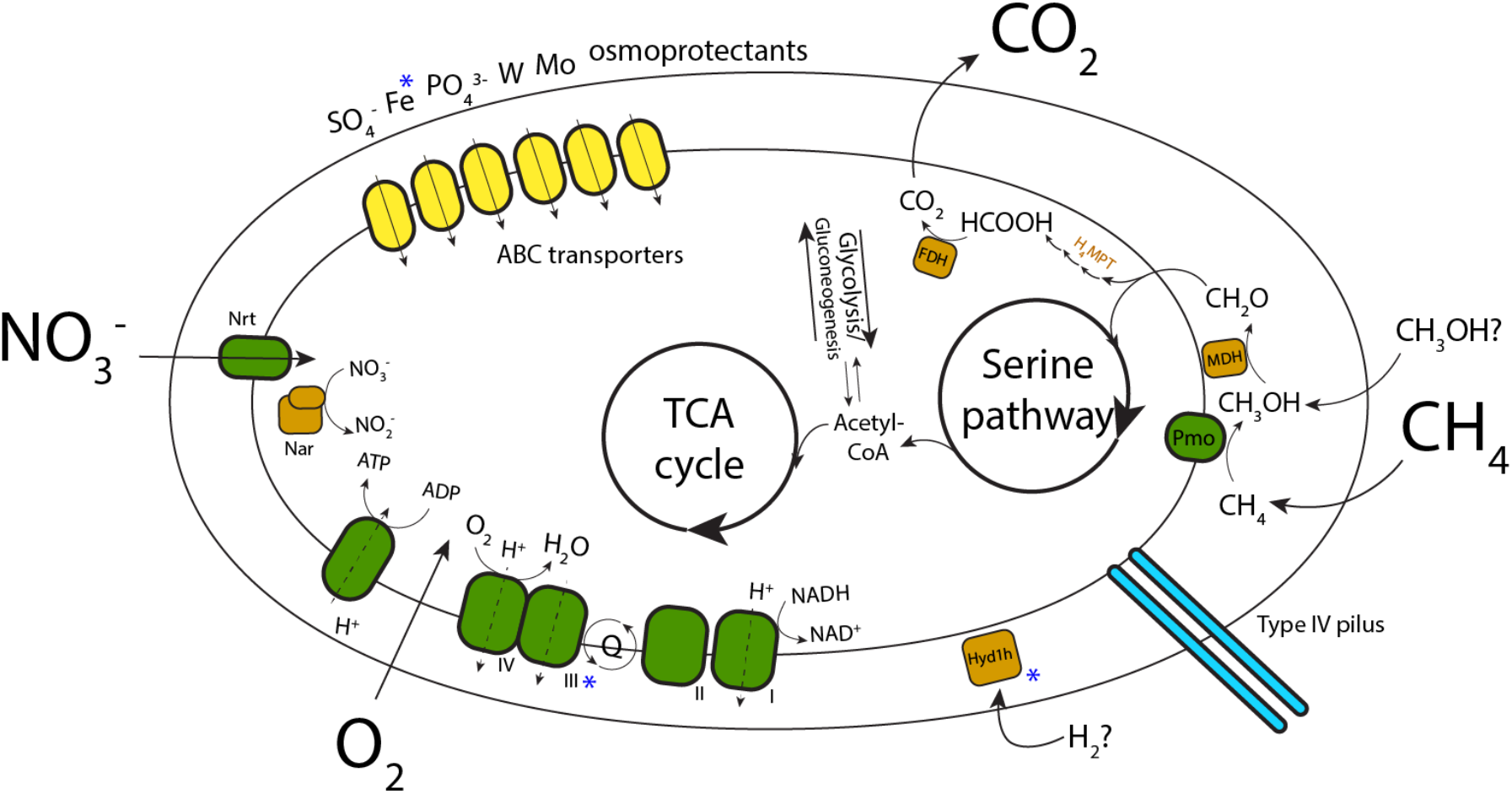
Conceptual schematic of the energy metabolism of wb1-P19 and its potential interactions with the cave environment, based on metagenomic reconstruction of the Lehman and Frasassi MAGs. Blue asterisks indicate functions encoded in the Frasassi but not the Lehman MAG.

USCg-Taylor is a member of the uncultivated upland soil cluster γ (USCγ) of high-affinity methanotrophs [19]. Consistent with this, the wb1-P19 MAGs from Lehman and Frasassi encode a particulate methane monooxygenase (*pmoCAB* operon) that is related to methane monooxygenases from USCγ methanotrophs, including representative USCγ sequences [20, 21] (Figure S2). They encode a complete methanotrophic pathway that includes enzymes for methanol, formaldehyde, and formate oxidation, along with all the genes for the serine pathway for carbon assimilation during methane or methanol oxidation. Neither of the wb1-P19 MAGs encode RuBisCO or any known CO_2_ fixation pathways, indicating that they are heterotrophs. Indeed, both bins encode complete TCA cycles and have major respiratory complexes I, II, and IV. In addition to encoding genes for oxygen-reducing terminal oxidases CoxAB and CydAB, they also encode a membrane-bound dissimilatory nitrate reductase (NarGHJI), indicating that they are facultative anaerobes capable of both oxygen and nitrate reduction. The wb1-P19 MAG from Frasassi is larger than the Lehman Caves MAG and encodes additional cbb3- and bo3-type terminal oxidases for oxygen reduction (CcoNO and CyoAD) and a group 1h hydrogenase that indicate more flexibility in substrates for energy catabolism. The USCg_Taylor, and JALZJW01 MAGs encode a urease that is absent from the Lehman or Frasassi MAGs, indicating diverse nutrient acquisition capabilities in this group beyond that represented in the two cave MAGs described here.

Previous studies have shown that USCγ methanotrophs are abundant in caves [3, 22-25], and the assignment of wb1-P19 to this group is further evidence for the ubiquity of these high-affinity methanotrophs in diverse cave ecosystems. Recently, the name “*Candidatus* Methyloligotrophales” was proposed for an order that includes USCγ [22], so while we don’t yet know if wb1-P19 represent all or just certain clades of USCγ organisms, rRNA genes classified as wb1-P19 should be considered part of this larger group. Given the importance of caves as global methane sinks, and the abundance of wb1-P19 in diverse karst and volcanic caves, these organisms are likely globally important for methane consumption in subterranean ecosystems worldwide. Many caves are energy-limited environments, so their abundance may reflect the importance of atmospheric methane in the absence of other substrates for microbial growth. We hope that the recognition of wb1-P19 as high-affinity methanotrophs helps lead to new culture-dependent and -independent analyses of their activity and ecophysiology in Earth’s subsurface.

## Acknowledgements

We thank the LD Hose, GM Baker, JD Powell, PF Dye, and the staff at Great Basin National Park for assisting with sample collection and permitting in Lehman Caves. Special thanks to A. Montanari for the use of facilities at the Osservatorio Geologico di Coldigioco, and to S. Recanatini for logistical and field support in Italy. Thanks to the staff at the University of Minnesota Genomics Center for insightful discussion and assistance with sequencing challenging samples, and to C Knief for expert advice on USCγ methanotrophs. Funding was provided by the National Cave and Karst Research Institute (NCKRI), the National Science Foundation (NSF EAR 2239710), the National Park Service (award P24AC00075-00), the United States Forest Service (award 24-CS-11041700-001), and the Rocky Mountain Association of Geologists (Gary L. Babcock Memorial Scholarship awarded to ZEH). Samples from Lehman Caves were collected under National Park Service permits GRBA-2018-SCI-0012 and GRBA-2019-SCI-0010.

## Notes

### Competing Interest Statement

BA is an employee of Phase Genomics, the developer of metagenomic proximity ligation technology.

## REFERENCES

1. Lennon JT, Nguyễn-Thùy D, Phạm TM et al. Microbial contributions to subterranean methane sinks. Geobiology. 2017;15:254–58 10.1111/gbi.12214

2. Waring CL, Hankin SI, Griffith DWT et al. Seasonal total methane depletion in limestone caves. Scientific Reports. 2017;7:8314 10.1038/s41598-017-07769-6

3. Webster Kevin D, Schimmelmann A, Drobniak A et al. Diversity and composition of methanotroph communities in caves. Microbiology Spectrum. 2022;10:e01566–21 10.1128/spectrum.01566-21

4. Fernandez-Cortes A, Cuezva S, Alvarez-Gallego M et al. Subterranean atmospheres may act as daily methane sinks. Nature Communications. 2015;6:7003 10.1038/ncomms8003

5. Martin JB. Carbonate minerals in the global carbon cycle. Chemical Geology. 2017;449:58–72 10.1016/j.chemgeo.2016.11.029

6. Martin-Pozas T, Cuezva S, Fernandez-Cortes A et al. Role of subterranean microbiota in the carbon cycle and greenhouse gas dynamics. Science of The Total Environment. 2022;831:154921 10.1016/j.scitotenv.2022.154921

7. Holmes AJ, Tujula NA, Holley M et al. Phylogenetic structure of unusual aquatic microbial formations in Nullarbor caves, Australia. Environmental Microbiology. 2001;3:256–64 10.1046/j.1462-2920.2001.00187.x

8. Yilmaz P, Parfrey LW, Yarza P et al. The SILVA and “All-species Living Tree Project (LTP)” taxonomic frameworks. Nucleic Acids Research. 2014;42:D643–D48 10.1093/nar/gkt1209

9. Hathaway JJM, Salazar-Hamm PS, Caimi NA et al. Comparison of fungal and bacterial microbiomes of bats and their cave roosting environments at El Malpais National Monument, New Mexico, USA. Geomicrobiology Journal. 2024;41:82–97 10.1080/01490451.2023.2283427

10. Jurado V, Gonzalez-Pimentel JL, Miller AZ et al. Microbial communities in vermiculation deposits from an alpine cave. Frontiers in Earth Science. 2020;8 10.3389/feart.2020.586248

11. Martin-Pozas T, Fernandez-Cortes A, Cuezva S et al. New insights into the structure, microbial diversity and ecology of yellow biofilms in a Paleolithic rock art cave (Pindal Cave, Asturias, Spain). Science of The Total Environment. 2023;897:165218 10.1016/j.scitotenv.2023.165218

12. Weng MM, Zaikova E, Millan M et al. Life underground: Investigating microbial communities and their biomarkers in Mars-analog lava tubes at Craters of the Moon National Monument and Preserve. Journal of Geophysical Research: Planets. 2022;127:e2022JE007268 10.1029/2022JE007268

13. Zhu H-Z, Zhang Z-F, Zhou N et al. Diversity, distribution and co-occurrence patterns of bacterial communities in a karst cave system. Frontiers in Microbiology. 2019;10 10.3389/fmicb.2019.01726

14. Ghezzi D, Jiménez-Morillo NT, Foschi L et al. The microbiota characterizing huge carbonatic moonmilk structures and its correlation with preserved organic matter. Environmental Microbiome. 2024;19:25 10.1186/s40793-024-00562-9

15. Havlena ZE, Hose LD, DuChene HR et al. Origin and modern microbial ecology of secondary mineral deposits in Lehman Caves, Great Basin National Park, NV, USA. Geobiology. 2024;22:e12594 10.1111/gbi.12594

16. Gogoleva N, Chervyatsova O, Balkin A et al. Microbial tapestry of the Shulgan-Tash cave (Southern Ural, Russia): influences of environmental factors on the taxonomic composition of the cave biofilms. Environmental Microbiome. 2023;18:82 10.1186/s40793-023-00538-1

17. Eren AM, Kiefl E, Shaiber A et al. Community-led, integrated, reproducible multi-omics with anvi’o. Nature Microbiology. 2021;6:3–6 10.1038/s41564-020-00834-3

18. Parks DH, Chuvochina M, Rinke C et al. GTDB: an ongoing census of bacterial and archaeal diversity through a phylogenetically consistent, rank normalized and complete genome-based taxonomy. Nucleic Acids Research. 2022;50:D785–D94 10.1093/nar/gkab776

19. Edwards Collin R, Onstott Tullis C, Miller Jennifer M et al. Draft genome sequence of uncultured upland soil cluster Gammaproteobacteria gives molecular insights into high-affinity methanotrophy. Genome Announcements. 2017;5:10.1128/genomea.00047-17 10.1128/genomea.00047-17

20. Knief C. Diversity and habitat preferences of cultivated and uncultivated aerobic methanotrophic bacteria evaluated based on pmoA as molecular marker. Frontiers in Microbiology. 2015;6 10.3389/fmicb.2015.01346

21. Knief C, Lipski A, Dunfield Peter F. Diversity and activity of methanotrophic bacteria in different upland soils. Applied and Environmental Microbiology. 2003;69:6703–14 10.1128/AEM.69.11.6703-6714.2003

22. Bay SK, Ni G, Lappan R et al. Microbial aerotrophy enables continuous primary production in diverse cave ecosystems. bioRxiv. 2024:2024.05.30.596735 10.1101/2024.05.30.596735

23. Cheng X, Wang H, Zeng Z et al. Niche differentiation of atmospheric methane-oxidizing bacteria and their community assembly in subsurface karst caves. Environmental Microbiology Reports. 2022;14:886–96 10.1111/1758-2229.13112

24. Cheng X-Y, Liu X-Y, Wang H-M et al. USCγ dominated community composition and cooccurrence network of methanotrophs and bacteria in subterranean karst caves. Microbiology Spectrum. 2021;9:10.1128/spectrum.00820-21 10.1128/spectrum.00820-21

25. Zhao R, Wang H, Cheng X et al. Upland soil cluster γ dominates the methanotroph communities in the karst Heshang Cave. FEMS Microbiology Ecology. 2018;94:fiy192 10.1093/femsec/fiy192

