## Supplementary Methods, Tables, and Figures for "Metagenomic insights into an enigmatic gammaproteobacterium that is important for carbon cycling in cave ecosystems worldwide"

### Supplementary Materials and Methods

#### Sample collection, DNA extraction, and library preparation for Lehman Caves samples

Samples were collected in 2018 and 2019 from Lehman Caves (39.0054 °N, 114.2207 °W) in Great Basin National Park, as described in Havlena et al. [1]. DNA was isolated from samples LMC19-11 and LGA18-4 using the DNeasy Powersoil Pro DNA Isolation Kit (Qiagen, Germantown, MD, USA) with the bead beating protocol that was shown by Havlena et al. [1] to maximize DNA recovery.

Metagenomic libraries were prepared from sample LMC19-11 using the NexteraXT library protocol, and from sample LGA18-4 using the Illumina DNA Prep (formerly Nextera DNA Flex) protocol. Libraries were sequenced on an Illumina NovaSeq S1 flowcell with 150 paired end cycles (Illumina, Inc., San Diego, CA, USA) at the University of Minnesota Genomics Center.

#### Sample collection, DNA extraction, and library preparation for Frasassi Caves samples

For the Frasassi Caves metagenome, samples of biovermiculations [2] were collected from Pozzo dei Cristalli in January, 2019. Samples were kept cool but not frozen for 3 days during transport to prevent cell lysis and DNA shearing during freeze-thaw cycles, and then subsequently stored and -80$^{o}$C until library preparation.

Proximity ligation libraries were prepared using the ProxiMeta Hi-C library preparation kit (Phase Genomics, Seattle, WA) using the manufacturer provided protocol. Shotgun and Hi-C libraries were sequenced on a NovaSeq S1 flowcell (Illumina). High-quality metagenome-assembled genomes were generated from the Hi-C data using ProxiMeta Proximity-Guided Assembly for a complex metagenome, using Phase Genomics' automated pipeline [3]. At this time, we are only describing one out of many bins generated from this process; the others (from this and additional samples) will be reported in an upcoming publication.

#### Metagenomic analysis

Quality filtering and trimming, assembly, and binning of the Lehman metagenomes were performed as in Jones et al. [4]. Briefly, reads were trimmed, filtered, and residual adaptors removed using sickle v. 1.33 (https://github.com/najoshi/sickle) and cutadapt v. 4.2 [5] to a final minimum length of 50 bp and mean quality score ≥ 28 (3' trimming only). Reads were assembled metaSPAdes v. 3.15.5 using k-mer sizes of 21, 33, and 55 [6], either using the full datasets or subsampled datasets in which either 20% or 50% of the quality trimmed and filtered reads were randomly selected. Read mapping was performed with bowtie2 v. 2.5.1 and samtools v. 1.6 [7, 8]. Contigs were binned with MetaBat2 using default parameters [9]. Bins were then manually evaluated with anvi'o v. 7.1 [10] (Supplementary Figure S1) and checked for completeness and contamination in anvi'o and CheckM and CheckM2 [11, 12]. We also compared binning strategies using MaxBin 2.0 [13], with DAS Tool [14] to combine assemblies, but ultimately received the best results in this case from MetaBat2 alone.

For gene searching in unbinned assemblies, protein-coding sequences were annotated with Prodigal v. 2.6.3 [15] in metagenomic gene calling mode, and for metagenome-assembled genomes, Prodigal was used in single genome mode. Protein-coding genes were annotated according to KEGG Orthology (KO) categories with GhostKOALA [16]. rRNA genes were identified using *barrnap* v. 0.9 (https://github.com/tseemann/barrnap), *rpoB* sequences were identified using HMMer (http://hmmer.org), and additional gene searching was performed with

BLAST [17]. We identified certain genes indicative of lithotrophic metabolisms using LithoGenie [18], and confirmed and classified hydrogenases annotated with Lithogenie with using HydDB [19].

We used BLASTP to identify a gene encoding hydroxypyruvate reductase (Hpr) that was missing from the KEGG annotations of the Lehman Caves MAG. To identify this missing gene, we downloaded some of the sequences from the Hpr phylogeny from But et al. [20], and used BLASTP to compare them against open reading frames. We identified a high-scoring *hpr* homolog on the same contig as other enzymes in the serine pathway that was evidently missed by the KEGG annotation, this confirming the complete serine pathway for carbon assimilation. We also used HMMer to search the Lehman metagenome for RuBisCO large subunit genes (*rbcL*) and ensure that there were no high-coverage genes in the dataset that were unbinned, and that the absence of these genes that encode enzymes for $CO_2$ fixation were truly absent from the metagenome.

**Phylogenetic analysis**

16S rRNA genes were aligned in ARB v. 6.0.4 [21], and positions with more than 50% gaps were removed prior to analysis for a final alignment length of 1333 positions. Maximum likelihood analysis was computed with RAxML version 8.2.12 [22] using the general time-reversible model with gamma distributed rates and invariable sites estimated from the data (model GTRGAMMAIX).

For phylogenetic analysis of RpoB and PmoA/AmoA, sequences were aligned using the Expresso algorithm in T-Coffee [23]. Analyses included the top 5 BLASTP matches to the Lehman Caves RpoB and PmoA, and in the case of PmoA, representative sequences of the USCg organisms from Knief [24]. Maximum likelihood analyses were computed with RAxML using the LG amino acid substitution model [25] with empirical frequencies, gamma distributed rates and invariable sites estimated from the data (model PROTGAMMAILGF).

### Supplementary Figures and Tables

**Table S1.** Summary of Lehman Caves and Frasassi wb1-P19 MAGs

| | Lehman Caves Bin3 (20% subsample) | Frasassi Caves bin PCtop12 |
|---|---|---|
| Genome size (Mbp) | 2.36 | 4.56 |
| Total scaffolds | 231 | 415 |
| N50 (Kbp) | 13.1 | 15.6 |
| Mean scaffold length (Kbp) | 10.2 | 11 |
| Longest scaffold (Kbp) | 57.4 | 58.5 |
| % G+C | 60.6 | 59.6 |
| Predicted genes | 2356 | 4774 |
| Completeness (%), anvi'o | 95.8 | |
| Redundancy (%), anvi'o | 0 | |
| Completeness (%), checkm) | 89.98 | 91.55 |
| Contamination (%), checkm | 0.34 | 4.12 |
| Strain heterogeneity (%), checkm | 0 | 39.13 |
| Completeness (%), checkm2 | 95.8 | 98.99 |
| Contamination (%), checkm2 | 0.02 | 3.25 |

**Table S2.** Select genomic capabilities of the two wb1-P19 metagenome-assembled genomes

| Description | Gene | KO annotations/other evidence | Lehman | Frasassi |
|---|---|---|---|---|
| **Central metabolism, nutrient acquisition** | | | | |
| Glycolysis/gluconeogenesis | | | Complete | Complete |
| TCA cycle | | | Complete | Complete |
| Type IV pilus | *pilABC* | K02650, K02652, K02653 | present | present |
| Sulfate assimilation | | | | |
| phosphoadenosine phosphosulfate reductase [EC:1.8.4.8 1.8.4.10] | *cysH* | K00390 | present | present |
| adenylylsulfate kinase [EC:2.7.1.25] | *cysC* | K00860 | present | present |
| sulfite reductase (ferredoxin) [EC:1.8.7.1] | *sir* | K00392 | present | present |
| sulfate adenylyltransferase [EC:2.7.7.4] | *sat* | K00958 | present | present |
| ABC transporters (KEGG annotations) | | | | |
| tungstate | *tupACB* | K05772, K06857, K05773 | present | present |
| molybdate | *modABCE* | K02020, K02018, K02017, K02019 | present | present |
| phosphate | *pstBACS* | K02036, K02038, K02037, K02040 | present | present |
| osmoprotectant | *opu* | K05845, K05846, K05847 | present | present |
| iron | *afuABC* | K02010, K02011, K02012 | - | present |
| **Oxygen reduction** | | | | |
| cytochrome bd ubiquinol oxidase | *cydAB* | K00425, K00426 | present | present |
| cytochrome c oxidase | *coxABCD* | K02275, K02274, K02276, K02277 | present | present |
| cytochrome c oxidase cbb3-type subunit I [EC:7.1.1.9] | *ccoNO* | K00404, K00405 | - | present |
| cytochrome o ubiquinol oxidase | *cyoABCD* | K02297, K02298, K02299, K02300 | - | present |
| **Nitrate reduction** | | | | |
| nitrate/nitrite transport system | *nrtBCD* | K15576, K15577, K15578, K15579 | *nrtBCD* only | *nrtABCD* |
| nitrate reductase / nitrite oxidoreductase | *narGHJI* | K00370, K00371, K00373, K00374 | present | present |
| nitrite reductase (NADH) | *nirBD* | K00362, K00363 | *nirB* only | *nirBD* |
| **Hydrogen oxidation** | | | | |
| Group 1h hydrogenase (Respiratory $H_2$-uptake [NiFe] hydrogenases) | | HydDB annotation | - | present |
| **Methane oxidation** | | | | |
| Methane oxidation | | | | |
| particulate methane monooxygenase | *pmoCAB* | K10946, K10944, K10945 | present | present |
| Methanol oxidation | | | | |
| methanol dehydrogenase (cytochrome c) (*mdh*) | *mdh12* | K14028. K14029 | present | present |
| methanol dehydrogenase, lanthanide-dependent (*xoxF*) | *xoxF* | K23995 | present | present |
| Formaldehyde oxidation (Tetrahydromethanopterin pathway) | | | | |
| 5,6,7,8-tetrahydromethanopterin hydro-lyase; formaldehyde-activating enzyme | *fae* | K10713 | present | present |
| methylene-tetrahydromethanopterin dehydrogenase | *mtdB* | K10714 | present | present |
| methenyltetrahydromethanopterin cyclohydrolase | *mch* | K01499 | present | present |
| formylmethanofuran--tetrahydromethanopterin N-formyltransferase | *ftr* | K00672 | present | present |
| formylmethanofuran dehydrogenase | *fwdABCD* | K00200, K00201, K00202, K00203 | present | present |
| Formate oxidation | | | | |
| formate dehydrogenase | FDH | K00122 | present | present |
| **Serine pathway for carbon assimilation** | | | | |
| glycine hydroxymethyltransferase | *glyA* | K00600 | present | present |
| serine-glyoxylate transaminase | *sga* | K00830 | present | present |
| hydroxypyruvate reductase | *hpr* | IDed by BLAST | present | present |
| glycerate 2-kinase | *glx* | K00865 | present | present |
| enolase | *eno* | K01689 | present | present |
| phosphoenolpyruvate carboxylase | *ppc* | K01595 | present | present |
| malate dehydrogenase | *mdh* | K00024 | present | present |
| malate-CoA ligase | *mtkAB* | K08692, K14067 | present | present |
| malyl-CoA lyase | *mcl* | K08691 | present | present |

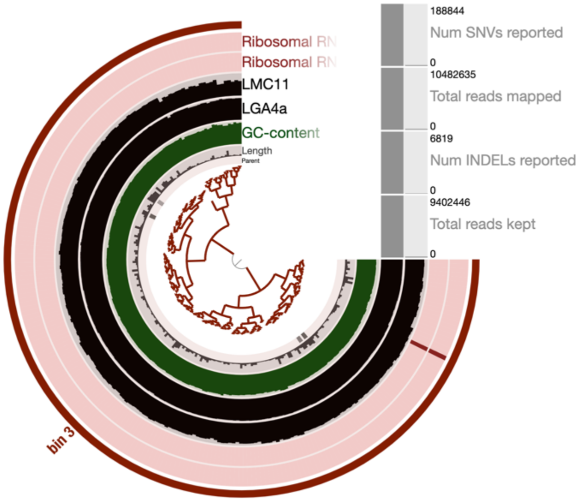


**Figure S1.** Screen captures showing wb1-P19 bins in anvi'o. The left image is of the bin generated by subsampling 20% of the dataset, and the right image is the bin from the full dataset. The red bars in the outer two pink circles show the location of the rRNA genes, and the two layers outside of the GC content bar graph show the average coverage of splits in the two Lehman metagenomes LGA18-4a and LMC19-11a, the latter of which was too small for binning. The bin generated following subsampling was slightly larger and more complete than the bin generated from the full dataset (estimated 95.8% complete and 2.36 Mb for the former versus 94.4% complete and 2.26 Mb for the latter; both 0% redundancy).

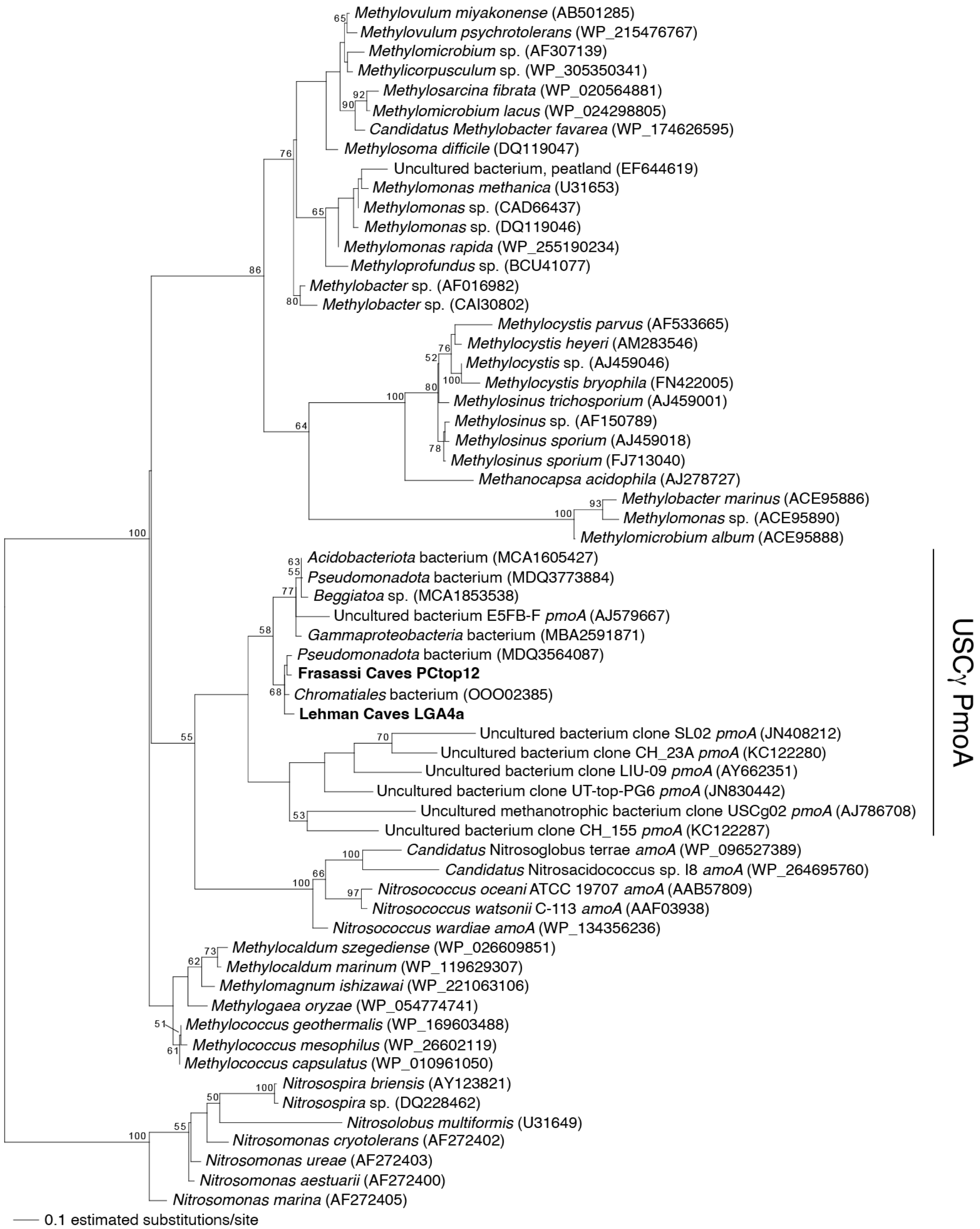


**Figure S2.** Maximum likelihood phylogenetic analysis of PmoA from the Lehman and Frasassi metagenomes. The phylogenetic analysis includes the top BLAST matches to the Lehman and Frasassi PmoA, as well as representative PmoA sequences from USCγ organisms from Knief [24], as well as AmoA from several members of the *Nitrosococcaceae*. Numbers on nodes indicate bootstrap support >50.

### REFERENCES


1. Havlena ZE, Hose LD, DuChene HR *et al.* Origin and modern microbial ecology of secondary mineral deposits in Lehman Caves, Great Basin National Park, NV, USA. *Geobiology*. 2024;**22**:e12594 https://doi.org/https://doi.org/10.1111/gbi.12594
2. Jones DS, Lyon EH, Macalady JL. Geomicrobiology of biovermiculations from the Frasassi cave system, Italy. *Journal of Cave and Karst Studies*. 2008;**70**:78-93
3. Burton JN, Liachko I, Dunham MJ *et al.* Species-level deconvolution of metagenome assemblies with Hi-C–based contact probability maps. *G3 Genes|Genomes|Genetics*. 2014;**4**:1339-46 https://doi.org/10.1534/g3.114.011825
4. Jones DS, Walker GM, Johnson NW *et al.* Molecular evidence for novel mercury methylating microorganisms in sulfate-impacted lakes. *The ISME Journal*. 2019;**13**:1659-75 https://doi.org/10.1038/s41396-019-0376-1
5. Martin M. Cutadapt removes adapter sequences from high-throughput sequencing reads. *EMBnetjournal; Vol 17, No 1: Next Generation Sequencing Data AnalysisDO - 1014806/ej171200*. 2011
6. Nurk S, Meleshko D, Korobeynikov A *et al.* metaSPAdes: a new versatile metagenomic assembler. *Genome Research*. 2017;**27**:824-34 https://doi.org/10.1101/gr.213959.116
7. Langmead B, Salzberg SL. Fast gapped-read alignment with Bowtie 2. *Nature Methods*. 2012;**9**:357-59 https://doi.org/10.1038/nmeth.1923
8. Danecek P, Bonfield JK, Liddle J *et al.* Twelve years of SAMtools and BCFtools. *GigaScience*. 2021;**10**:giab008 https://doi.org/10.1093/gigascience/giab008
9. Kang DD, Li F, Kirton E *et al.* MetaBAT 2: an adaptive binning algorithm for robust and efficient genome reconstruction from metagenome assemblies. *PeerJ*. 2019;**7**:e7359 https://doi.org/https://doi.org/10.7717/peerj.7359
10. Eren AM, Esen ÖC, Quince C *et al.* Anvi'o: an advanced analysis and visualization platform for 'omics data. *PeerJ*. 2015;**3**:e1319
11. Parks DH, Imelfort M, Skennerton CT *et al.* CheckM: assessing the quality of microbial genomes recovered from isolates, single cells, and metagenomes. *Genome Research*. 2015;**25**:1043-55 https://doi.org/10.1101/gr.186072.114
12. Chklovski A, Parks DH, Woodcroft BJ *et al.* CheckM2: a rapid, scalable and accurate tool for assessing microbial genome quality using machine learning. *Nature Methods*. 2023;**20**:1203-12 https://doi.org/10.1038/s41592-023-01940-w
13. Wu Y-W, Simmons BA, Singer SW. MaxBin 2.0: an automated binning algorithm to recover genomes from multiple metagenomic datasets. *Bioinformatics*. 2016;**32**:605-07 https://doi.org/10.1093/bioinformatics/btv638
14. Sieber CMK, Probst AJ, Sharrar A *et al.* Recovery of genomes from metagenomes via a dereplication, aggregation and scoring strategy. *Nature Microbiology*. 2018;**3**:836-43 https://doi.org/10.1038/s41564-018-0171-1
15. Hyatt D, Chen G-L, LoCascio PF *et al.* Prodigal: prokaryotic gene recognition and translation initiation site identification. *BMC Bioinformatics*. 2010;**11**:119 https://doi.org/10.1186/1471-2105-11-119
16. Kanehisa M, Sato Y, Morishima K. BlastKOALA and GhostKOALA: KEGG tools for functional characterization of genome and metagenome sequences. *Journal of Molecular Biology*. 2016;**428**:726-31 https://doi.org/https://doi.org/10.1016/j.jmb.2015.11.006

17. Altschul SF, Madden TL, Schäffer AA *et al.* Gapped BLAST and PSI-BLAST: a new generation of protein database search programs. *Nucleic Acids Research*. 1997;**25**:3389-402 https://doi.org/10.1093/nar/25.17.3389
18. Garber A, Ramirez G, Merino N *et al.* MagicLamp: toolkit for annotation of genomic data using discreet and curated HMM sets. 2023: MagicLamp, GitHub repository:. https://github.com/Arkadiy-Garber/MagicLamp
19. Søndergaard D, Pedersen CNS, Greening C. HydDB: A web tool for hydrogenase classification and analysis. *Scientific Reports*. 2016;**6**:34212 https://doi.org/10.1038/srep34212
20. But SY, Egorova SV, Khmelenina VN *et al.* Biochemical properties and phylogeny of hydroxypyruvate reductases from methanotrophic bacteria with different c1-assimilation pathways. *Biochemistry (Moscow)*. 2017;**82**:1295-303 https://doi.org/10.1134/S0006297917110074
21. Ludwig W, Strunk O, Westram R *et al.* ARB: a software environment for sequence data. *Nucleic Acids Research*. 2004;**32**:1363-71 https://doi.org/10.1093/nar/gkh293
22. Stamatakis A. RAxML-VI-HPC: maximum likelihood-based phylogenetic analyses with thousands of taxa and mixed models. *Bioinformatics*. 2006;**22**:2688-90 https://doi.org/10.1093/bioinformatics/btl446
23. Armougom F, Moretti S, Poirot O *et al.* Expresso: automatic incorporation of structural information in multiple sequence alignments using 3D-Coffee. *Nucleic Acids Research*. 2006;**34**:W604-W08 https://doi.org/10.1093/nar/gkl092
24. Knief C. Diversity and habitat preferences of cultivated and uncultivated aerobic methanotrophic bacteria evaluated based on *pmoA* as molecular marker. *Frontiers in Microbiology*. 2015;**6** https://doi.org/10.3389/fmicb.2015.01346
25. Le SQ, Gascuel O. An improved general amino acid replacement matrix. *Molecular Biology and Evolution*. 2008;**25**:1307-20 https://doi.org/10.1093/molbev/msn067
